# Species activity promote the stability of fruit-frugivore interactions across a five-year multilayer network

**DOI:** 10.1101/421941

**Authors:** José M. Costa, Jaime A. Ramos, Sérgio Timóteo, Luís P. da Silva, Ricardo S. Ceia, Ruben H. Heleno

**Author notes:** CIBIO-InBIO, Research Centre in Biodiversity and Genetic Resources, University of Porto, Portugal (Luís P. da Silva).

## Abstract

Although biological communities are intrinsically dynamic, with both, species and interactions changing over time, interaction networks analyses to date are still largely static. We implemented a temporal multilayer network approach to explore the changes on species roles and on the emergent structure of a seed-dispersal network over five years. Network topology was relatively constant, with four well defined interaction modules spanning across all years. Importantly, species that were present on more years, were also disproportionally important on each year, thus forming a core of temporally reliable species that are critical to the cohesiveness of the multilayer network structure. We propose a new descriptor termed *species activity* that reflects the number of temporal, spatial or functional layers (e.g., different years, habitats, or functions) that each species integrates, providing a simple and powerful index of species importance for multilayer network cohesion.

## Introduction

The structure of ecological communities, reflected in the complex network of biotic interactions that connects all organisms and species, is intrinsically dynamic. Such dynamics may directly emerge from temporal changes on species composition (species turnover), switch in animal feeding preferences (rewiring), relative species abundances, and availability of suitable resources (phenological matching), such as flowers and fruits (Olesen *et al*. 2010; Burkle & Alarcón 2011; Trøjelsgaard & Olesen 2016). Although the dynamic nature of species interactions is widely recognized (Olesen *et al*. 2010; Trøjelsgaard & Olesen 2016), most studied networks to date are constrained to relatively short time scales. Several studies started to explore the temporal variability of species interaction networks across seasons and years, mostly focusing on plant-pollinator (Petanidou *et al*. 2008; Dupont *et al*. 2009; Chacoff *et al*. 2018) or on plant-frugivore interactions (Carnicer *et al*. 2009; González-Castro *et al*. 2012; Ramos-Robles *et al*. 2016). Nevertheless, although these studies provide useful information about structural community changes across time, they still inevitably aggregate all observed interactions into a few predefined and formally disconnected time-slices (e.g., years). Accordingly, they look for trends across collections of static interaction matrices, likely providing an incomplete perception of true temporal dynamics (Blonder *et al*. 2012; Pilosof *et al*. 2017). The tool to circumvent this limitation have recently became available, particularly with the implementation of a multilayer network approach where interdependencies between time-ordered layers (i.e., networks) are explicitly incorporated in the analysis by quantifying the strength of interlayer links that connect the same nodes (species) across multiple layers (Pilosof *et al*. 2017; García-Callejas *et al*. 2018; Timóteo *et al*. 2018). By independently quantifying intra- and interlayer strength, multilayer networks are a most powerful tool to explore meta-community dynamics, improving the realism of species interaction networks naturally spanning across multiple spatial (e.g., patches, habitats), temporal (e.g., seasons, years) or functional (e.g., herbivory, predation, parasitism) layers (Pilosof *et al*. 2017; Timóteo *et al*. 2018).

For most plants, seed-dispersal represents a key life-history stage through which they can colonize new habitats away from mother plants (Traveset *et al*. 2014). Birds are critical seed dispersers across most terrestrial ecosystems, largely influencing habitat structure and longterm vegetation dynamics (Jordano 2014; Wenny *et al*. 2016). Over the recent decades our understanding of the organization of plant-frugivore interaction networks has improved tremendously (Jordano *et al*. 2007; Donatti *et al*. 2011). Several studies monitored frugivory across multiple seasons (Carnicer *et al*. 2009; González-Castro *et al*. 2012; Ramos-Robles *et al*. 2016), usually revealing peaks in seed-dispersal intensity. In Southern Europe, this peak occurs during late-summer and early-autumn, where the production of fleshy fruits coincides with the post-breeding bird migration (Herrera 1984; Costa *et al*. 2018). Although both processes (fruit production and bird migration) occur every year, allowing the regular encounter between fruits and dispersers, there might be important fluctuations in their abundance and diversity (Herrera 1998). Surprisingly, we still do not know how these fluctuations affect the persistence of plant-frugivore interactions across years, hindering current understanding of long-term community dynamics (Estes *et al*. 2018). Furthermore, the lack of such a long-term perspective is unanimously recognized as a major limitation of current understanding of biodiversity – ecosystem services relationships as a whole (Tilman *et al*. 2014).

Here, we implemented a temporal multilayer network approach to explore the variability of seed-dispersal interactions across five consecutive years. Specifically, we aimed to (1) characterize and identify the drivers of species and interactions turnover; (2) investigate the relationships between short- and long-term species topological roles; and (3) explore the drivers of temporal changes in emergent network properties.

## Material and methods

### Experimental approach

For five consecutive years, we reconstructed the network of interactions between birds and fleshy-fruited plants on a secondary native forest in Central Portugal (40^<u>o</u>^19’N; 8^<u>o</u>^24’W). The site is under a typical Mediterranean climate and is dominated by *Quercus faginea*, *Arbutus unedo*, and *Pinus pinaster* with a dense and diverse understory dominated by native fleshy-fruited shrubs, such as *Pistacia lentiscus*, *Crataegus monogyna*, *Rhamnus alaternus*, *Rubus ulmifolius*, and *Viburnum tinus* (see detailed description in da Silva *et al*. 2016).

Interaction networks were assembled by identifying entire seeds on the droppings of mist-netted birds captured under two complementary designs: (1) sampling all days with favourable weather conditions during the month of September between 2012 and 2016 (corresponding to the predicted peak of seed-dispersal); and (2) sampling fortnightly between January and December 2013 to evaluate the intra-annual richness of pairwise seed-bird interactions. On each day, birds were captured with mist-nets (total length = 102 m), operated during the first five hours after dawn, and placed in individual cotton bags until they defecate or up to 30 minutes. All droppings retrieved from the bags were air-dried and the undamaged seeds were later extracted, counted and identified under a dissecting microscope with the aid of a seed reference collection. Interaction frequency was quantified as the number of droppings of bird species *i* containing undamaged seeds of plant species *j* (Vázquez *et al*. 2005; Heleno *et al*. 2013). This measure was used because it is more likely to reflect recruitment probability after post-dispersal density-dependent mortality of seeds deposited in the same dropping (Correia *et al*. 2016). The overall effort resulted in 635 sampling-hours distributed along 25, 17, 20, 21, and 20 days in September 2012 to 2016, respectively, and 120 hours in 24 additional days across the entire year of 2013. Sampling completeness was estimated for each year as the proportion of plant and bird species observed relatively to those estimated by the Chao2 richness estimator (Chao 1987) implemented in program EstimateS 9.1 (Colwell 2013). Additionally, fleshy-fruit availability was estimated by counting all ripe standing fruits along three linear transects (each: 25m × 2m) running parallel to the mist-nets and resampled each year in early-, mid-, and late-September. Unless stated otherwise, all results are presented as mean ± standard deviation.

### Interannual turnover of species and links

The interannual turnover of bird and plant species was assessed with the Whittaker beta diversity index (β_W_) (Whittaker 1960) adapted by Koleff *et al*. (2003), which varies between 0 (similar) and 1 (completely dissimilar species composition). The turnover of links was evaluated with package betalink (Poisot 2016) in R (R Core Team 2017), following the approach outlined in Poisot *et al*. (2012), which allows the partition of link turnover (β_WN_) between networks into two driving mechanisms: those attributed exclusively to species turnover (β_ST_) and those attributed to the rewiring of interactions between co-occurring species (β_OS_).

### Relationship between intra- and interannual species topological roles

To characterize the regularity of species across the five years, we propose a new species-level index in the context of ecological multilayer networks, which we coined “species activity”, and quantifies the number of layers (here: years) in which each species interacts (i.e., the number of layers with activity of each species). This descriptor is a direct extension of the concept of “node activity” used in physics to reflect the number of layers where the nodes of multiplex networks are active (Nicosia & Latora 2015). We then evaluated how species activity is related with species topological importance in each year, by computing three monolayer species-level descriptors: (1) degree, i.e., the number of mutualistic partners; (2) species strength, an estimation of the cumulative importance of each species for the species on the other trophic level (Barrat *et al*. 2004); and (3) specialization d’, quantifying species selectivity in relation to resource availability (Blüthgen *et al*. 2006). Additionally, we also evaluated the relationship between species activity and species versatility, a descriptor of multilayer centrality, expressing the sum of the importance of the partners of species *i*, both within and between layers (De Domenico *et al*. 2015b; Timóteo *et al*. 2018). Species versatility was computed using the PageRank algorithm (Brin & Page 2012) adapted to a multilayer scenario (De Domenico *et al*. 2015b) and available in program muxViz (De Domenico *et al*. 2015a). This was done separately for bird and plant species, based on unipartite projections of the original networks using the Newman’s method (Newman 2001) adapted for weighted networks (Opsahl 2013) with the R package tnet (Opsahl 2009).

In order to assess if plants are dispersed proportionally to their abundance on each year, we calculated the Kendall’s tau rank correlation test available from the R package Kendall (McLeod 2011), between the abundance of fleshy-fruits of each species in the transects and their respective interaction frequency. The effect of species activity on mean species degree and strength was assessed with generalized linear mixed models (GLMM) with Poisson and Gamma distributed errors, respectively. In order to control for the effect of variable network sizes, the number of species on the other trophic level (i.e., number of plant species for bird degree and vice-versa) was included as an offset variable in the Poisson GLMM. The relationship between species activity and species specialization d’ was modelled with linear mixed models (LMM). All mixed models were fitted with the R package lmer4 (Bates *et al*. 2015) and included year as a random factor. The relationship between species versatility and species activity was assessed with generalized linear models (GLM) with Gamma distributed errors.

### Interannual community structure

Changes in the emergent structure of the seed-dispersal network were evaluated by calculating four key network-level descriptors: (1) connectance,, the proportion of observed/ possible links (Jordano 1987); (2) network specialization H_2_’, measuring the community-level selectiveness of the observed interactions as a departure from a random (i.e., abundance-based) association pattern (Blüthgen *et al*. 2006); (3) weighted-interaction nestedness (WIN) (Galeano *et al*. 2009), quantifying how interactions are hierarchically organized (i.e., nested) around a core of the most generalist species (Bascompte *et al*. 2003); and (4) modularity, which identifies and quantifies the existence of groups of tightly interacting species, loosely linked to the remaining network (Olesen *et al*. 2007). To compute modularity, we employed an explicit multilayer approach where we included interlayer links connecting the same species occurring in consecutive years. These links were quantified as the change in each species relative abundance between consecutive layers (i.e., abundance *i* _t+1_ / abundance *i* _t_) (see Pilosof *et al*. 2017), where bird abundances correspond to the mean number of birds captured, and plant abundances corresponds to mean fruit availability in the transects. When plant species were found in the bird droppings but not in transects, these were attributed the lowest availability score (i.e., 1 fruit/transect), under the rationale that those fruits need to be available in order to be consumed but are probably locally rare. Modularity was maximized with a generalized Louvain algorithm (Blondel *et al*. 2008), implemented in MATLAB (The MathWorks, Inc., Natick, Massachusetts, USA) using code provided in Jutla *et al*. (2014) and modified by Pilosof *et al*. (2017) to account for the bipartite nature of the multilayer network (see also Timóteo *et al*. 2018). The significance of each descriptor was then assessed by comparing it with those obtained for 1000 randomized networks generated by a null model based on the Patefield’s algorithm (Patefield 1981), which randomly reshuffles the interactions across the matrix while constraining marginal totals. Each descriptor was considered significantly different from a random expectation if the respective z-score was lower than −1.96 or higher than 1.96, corresponding to a significance level of 0.05 (Trøjelsgaard *et al*. 2015). The randomized networks to compute modularity significance were obtained with the R package vegan (Oksanen *et al*. 2015). All other network-level descriptors and respective null-models were obtained with package bipartite (Dormann *et al*. 2008).

## Results

Throughout 2013 (fortnight sampling) we captured 671 birds from 30 species, whose 202 droppings contained 537 undamaged seeds from 16 plant species. September was the month with a greater diversity of links between fleshy-fruited plants and birds, with 15 out of the 40 links being detected in this month (Fig. 1).

**Figure 1.**
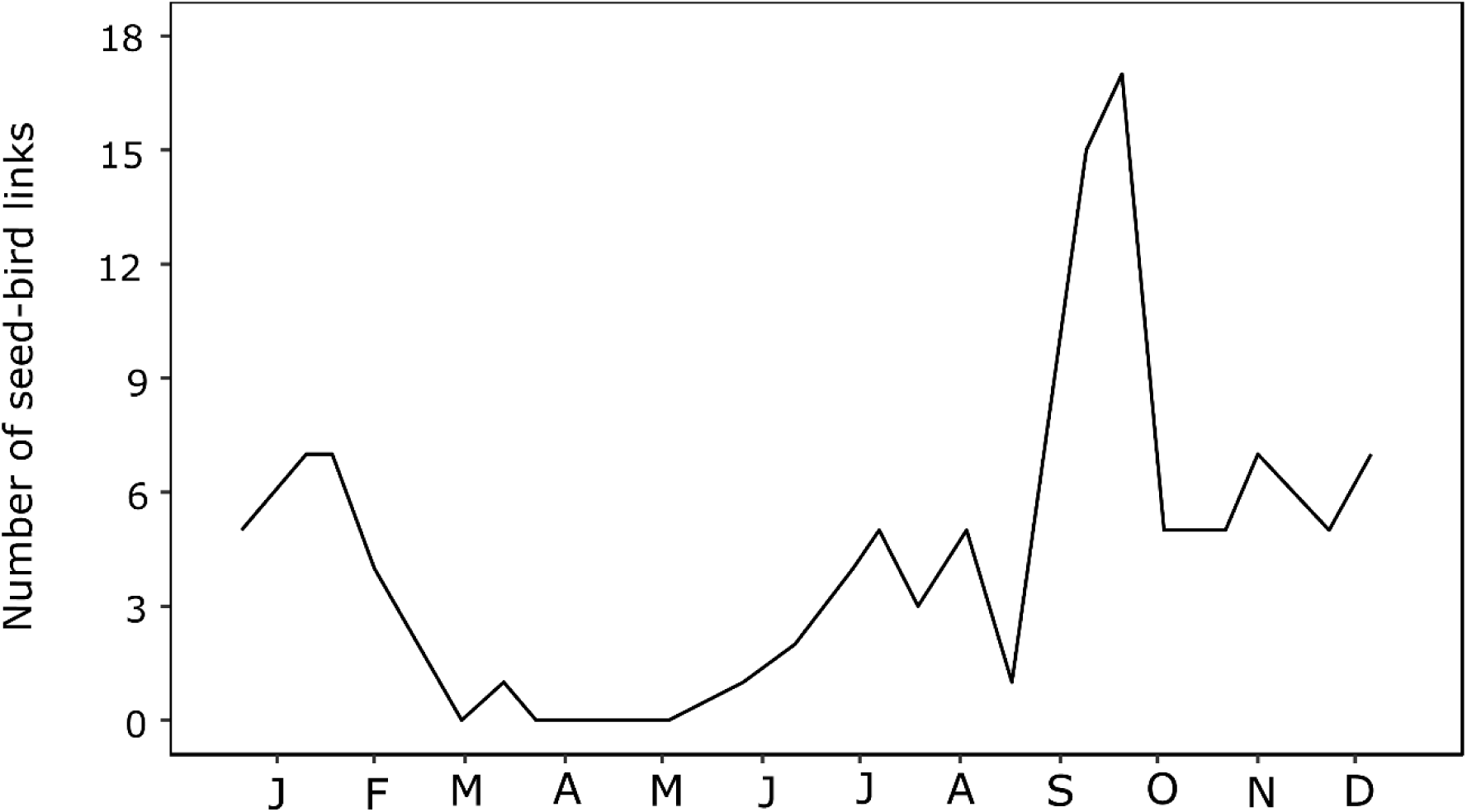
– Richness of pairwise links between seeds and their bird dispersers, recorded fortnightly throughout one year (2013) on a native forest in central Portugal.

Overall, the daily sampling throughout September from 2012 to 2016 resulted in the capture of 1620 birds (30 species), of which 454 (12 species) dispersed 2133 undamaged seeds from 17 plant species, rendering a total of 75 links (Fig. 2). Estimated sampling completeness was very high for both plants and birds, with an annual mean of 93% (Min.= 90%; Max.= 98%) and 92% (Min.= 89%; Max.= 100%) of species detected, respectively.

**Figure 2.**
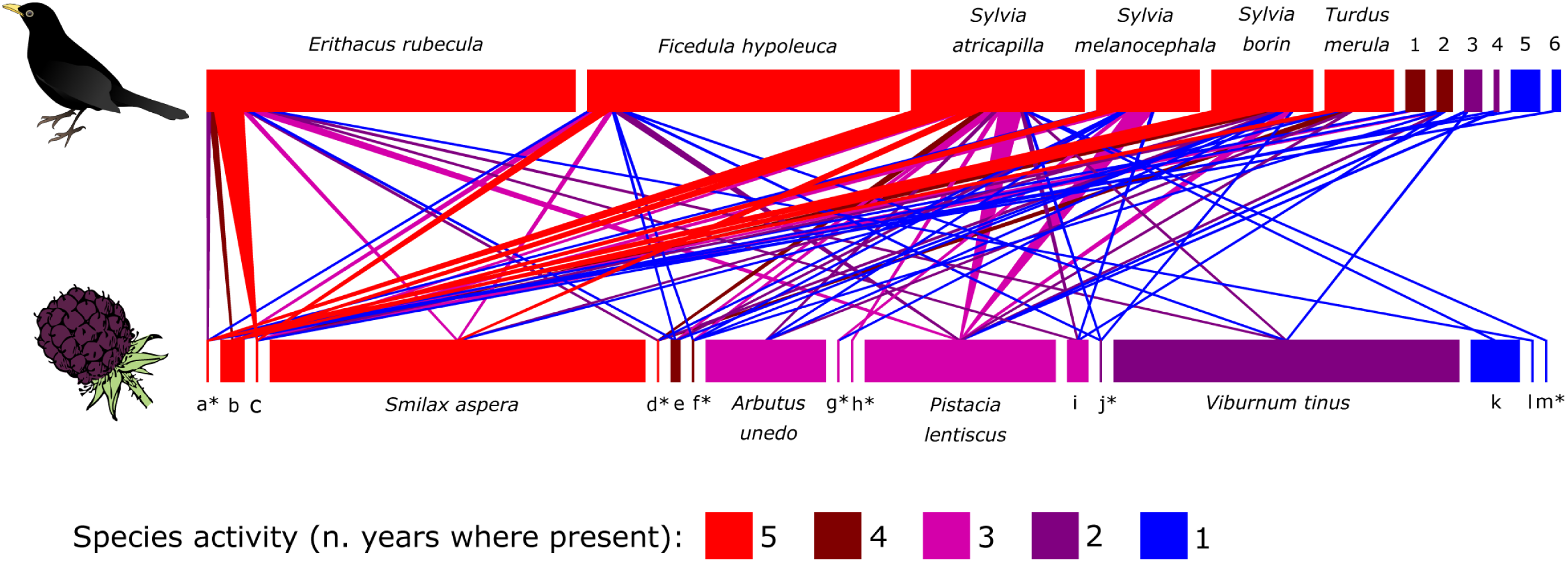
– Overall seed-dispersal network recorded during the peak of the fruiting season (September) across five consecutive years on a secondary native forest in central Portugal. Species are ordered from highest to lowest species activity, i.e. number of years where present. The width of the boxes representing plant and bird species is proportional to the number of fruits counted along linear transects and to the number of birds captured with mist-nets, respectively. Asterisks (^∗^) represent seed species recovered from bird droppings but not detected in the transects. 1 – *Cyanistes caeruleus*, 2 – *S. communis*, 3 – *Muscicapa striata*, 4 – *S. undata*, 5 – *Chloris chloris*, 6 – *Dendrocopos major;* Plants: a – *Ficus carica*, b – *Rhamnus alaternus*, c – *Rubus ulmifolius*, d – *Vitis vinifera*, e – *Phillyrea latifolia*, f – *Solanum nigrum*, g – *Daphne gnidium*, h – *Lonicera periclymenum*, i – *Rubia peregrina*, j – *Phytolacca americana*, k – *Crataegus monogyna*, l – *Olea europaea*, m – *Phillyrea angustifolia*.

### Interannual turnover of species and links

Five plant species (*Ficus carica*, *Rhamnus alaternus*, *Rubus ulmifolius*, *Smilax aspera*, and *Vitis vinifera*) and six bird species (*Erithacus rubecula*, *Ficedula hypoleuca*, *Sylvia atricapilla*, *S. borin*, *S. melanocephala*, and *Turdus merula*) were detected on the five years of the study (Figs. 2 and 3). These species accounted respectively for 29% of the fruit production and 50% of the individual birds captured in September across the five years. Mean species turnover between consecutive years (β_W_) was higher for plants than for birds (0.31 ± 0.12 and 0.16 ± 0.07, respectively).

Nine out of the 75 links detected (12%) were observed in all years, accounting on average for 49% (Min.= 30%; Max.= 63%) of the interactions detected in each year. The turnover of links was greater than that of plant and animal species (β_WN_= 0.53 ± 0.10) and mostly driven by interaction rewiring, i.e., by the detection of new links between species already co-occurring on previous years (β_OS_/β_WN_= 70% ± 14%) with a lower contribution of species turnover (β_ST_/β_WN_= 30% ± 14%).

### Relationship between intra- and interannual species topological roles

There was no significant correlation between fruit abundance and the frequency of interactions in any year (τ_2012_= 0.12, p= 0.74; τ_2013_= 0.60, p= 0.18; τ_2014_= 0.53, p= 0.07; τ_2015_= 0.44, p= 0.17; τ_2016_= −0.32, p= 0.63). Both plant and bird species activity were positively related to their respective mean degree (β_plants_±SE= 0.37 ± 0.09, χ^2^= 14.76, p < 0.01; β_birds_±SE= 0.68 ± 0.14, χ^2^= 24.45, p < 0.01; Fig. 3), mean species strength (β_plants_= 0.23 ± 0.04, χ^2^= 40.17, p < 0.01; β_birds_= 0.94 ± 0.11, χ^2^= 75.56, p < 0.01; Fig. 3), and versatility (β_plants_= 0.41 ± 0.06, χ^2^= 52.77, p < 0.01; β_birds_= −1.62 ± 0.18, χ^2^= 78.81, p < 0.01; Fig. 3). In contrast, plant and bird specialization d’ were not associated with species activity (β_plants_= 0.04 ± 0.02, χ^2^= 3.68, p= 0.06; β_birds_= 0.01 ± 0.02, χ^2^= 0.19, p= 0.66; Fig. 3).

**Figure 3.**
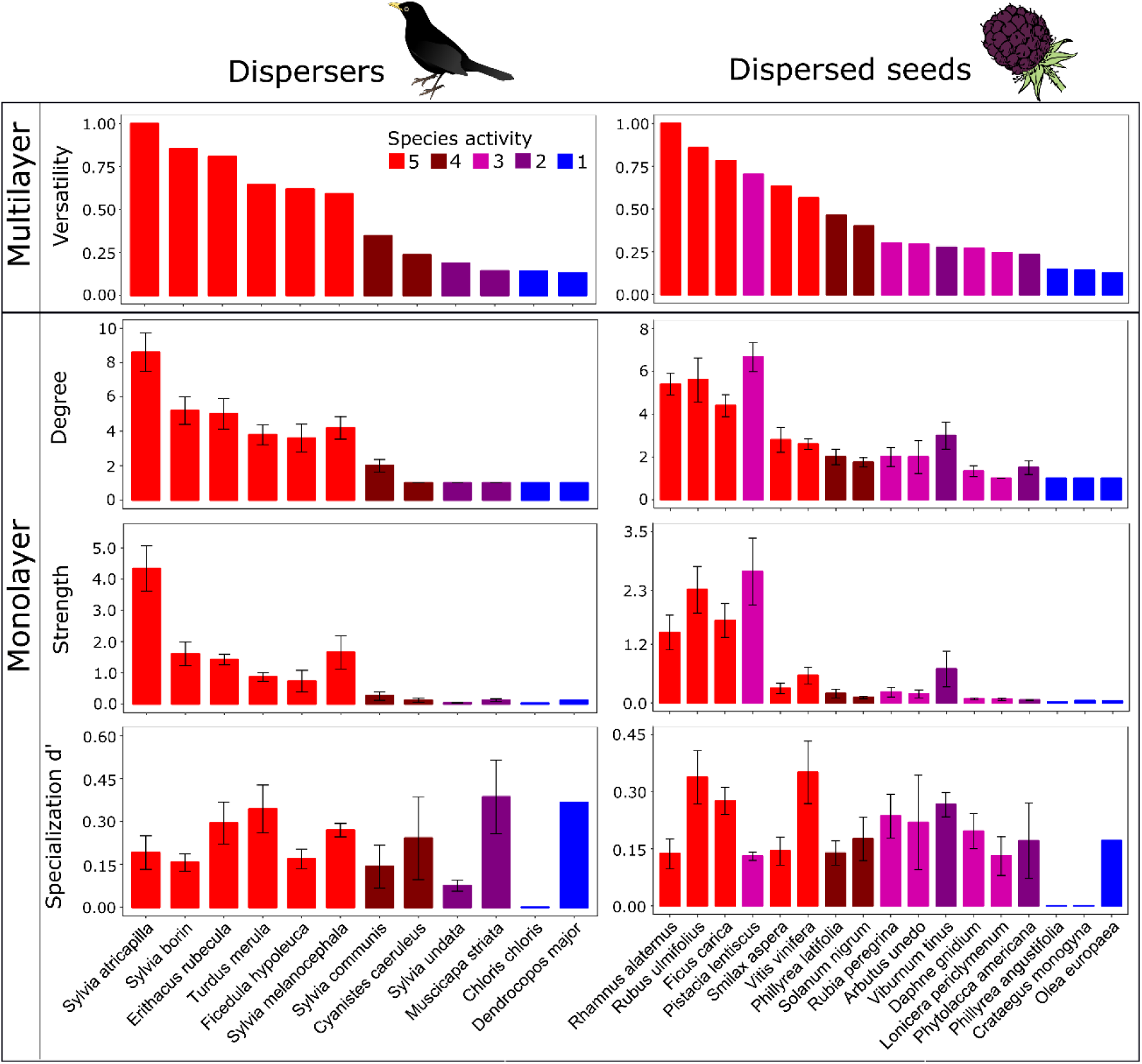
– Topological descriptors of species roles on avian seed-dispersal networks compiled over five years. The top panel corresponds to species roles on a temporal multilayer network, while the monolayer panels reflect average species roles across the yearly networks where each species occurs. Error bars represent the standard error for each descriptor across the five years. Bars without error bars correspond to species with no interannual variation to a given descriptor. Species are ordered according to their multilayer versatility (top).

### Interannual community structure

Overall, the topology of the networks was relatively constant throughout the study (Fig. 4). The network was significantly less connected (z_2012_= −5.12; z_2013_= −2.97; z_2014_= −2.15; z_2015_= −3.31; z_2016_= −4.51) and more specialized (z_2012_= 9.67; z_2013_= 3.90; z_2014_= 3.16; z_2015_= 5.86; z_2016_= 8.14) than predicted by the null models in all years. The network also tended to be significantly nested, which happened in all years except in 2014, when observed nestedness was indistinguishable from a random interaction pattern (z_2012_= 4.22; z_2013_= 3.43; z_2014_= 0.93; z_2015_= 3.91; z_2016_= 5.41).

**Figure 4.**
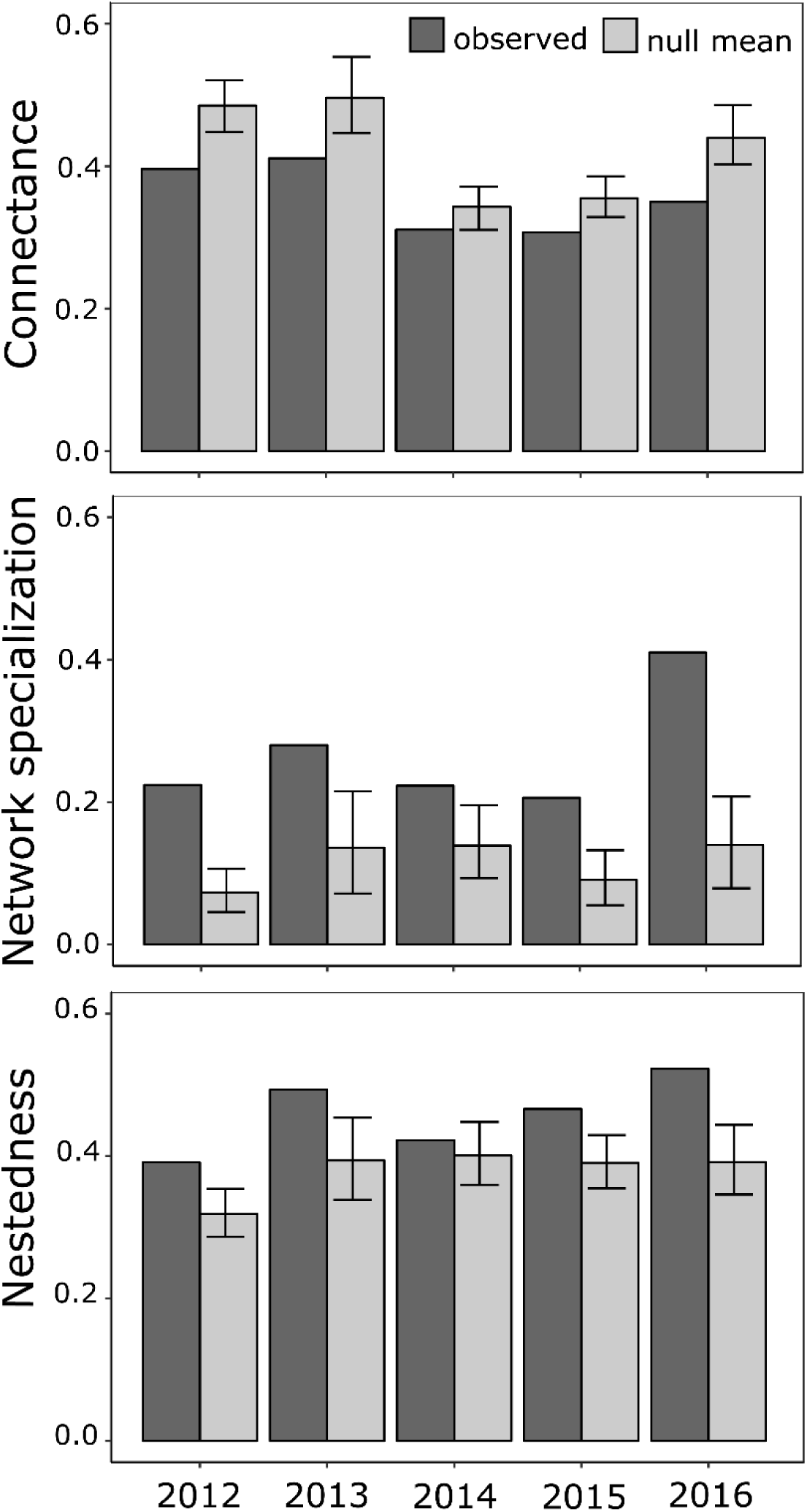
– Interannual variability in seed-dispersal network structure, namely: Connectance, Network specialization (H_2_’), and Nestedness (WIN). The observed descriptor (“observed”) is compared with the mean of 1000 Patefield null models (“null mean”). Error bars correspond to the 95% percentile of the null models’ distribution.

The overall multilayer network was significantly more modular than expected by chance (Q_obs_= 0.50, mean Q_null_= 0.43, z= 10.3), and formed by four interaction modules (Fig. 5) that spanned across the five years of the study. Most bird species (8 out of 10 species, 80%) were consistently allocated into the same module across all years. Plants had a lower temporal constancy regarding their module affiliation, with 9 out of the 14 plant species (64%) remaining in the same module across all years.

**Figure 5.**
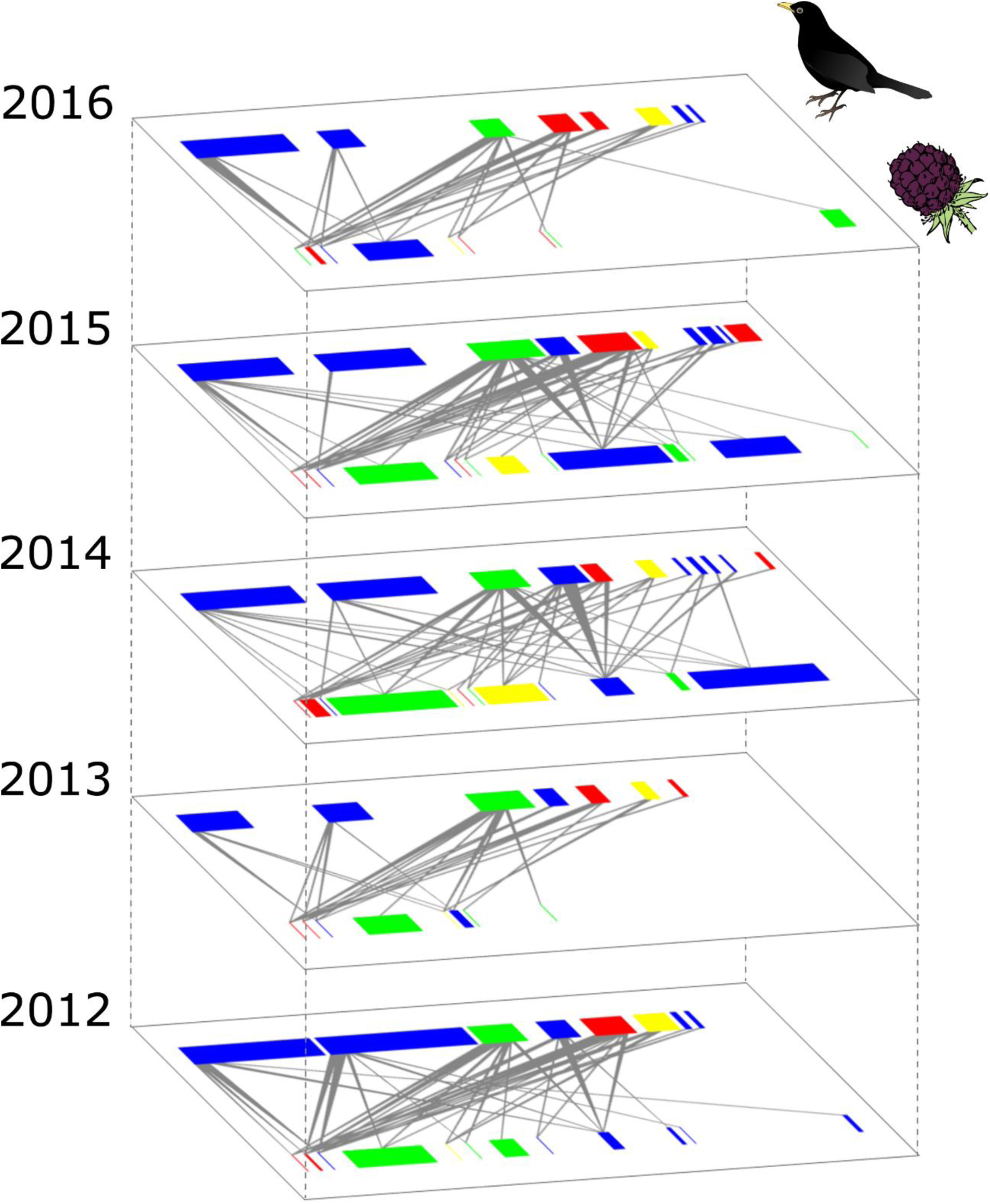
– Interannual module affiliation of species across a five-year temporal multilayer seed dispersal network (colours represents different interaction modules, see Methods). All networks are represented on the same scale and species are ordered as in figure 2. The width of the boxes representing plant and bird species is proportional to the number of fruits counted along linear transects and to the number of birds captured with mist nets, respectively.

## Discussion

Here, we show that the diversity of seed-dispersal interactions between birds and fleshy-fruited plants in Portugal is particularly high in September, when a peak in fruit production coincides with the passage of most migratory bird species. We compiled seed-dispersal interactions during this activity peak for five consecutive years to reconstruct a temporal multilayer network, quantifying intra- and interlayer (i.e., year) link strength. We found that the species present in more years (here said to have a greater species activity) also tend to be more important in each year, independently of their abundance. The emergent structure of the seed-dispersal network was very stable through time and included four well-defined interaction modules spanning across all years of the study. Altogether, our results suggest that the temporally most reliable species, which are not necessarily the most abundant, tend to occupy central roles in the seed-dispersal network across multiple temporal scales, thus providing a mechanism for network stability and increasing the reliability of the seed-dispersal service.

Species activity is a species-level multilayer descriptor that quantifies the role of each species as connectors in the multilayer network system, although it can also be used in a monolayer context (i.e., when inter-layer link strength is undefined). For example, this index has been implicitly used to explore the persistence of species across years in flower-visitor networks (Petanidou *et al*. 2008; Olesen *et al*. 2011b), across months in bird-fruit networks (Yang *et al*. 2013; Ramos-Robles *et al*. 2016), and across multiple habitat layers in seed dispersal networks (Timóteo *et al*. 2018). However, until now it was not properly described. This simple descriptor, naturally related to what Olesen *et al*. (2011b) named “temporal persistence”, is very intuitive and might prove useful in situations where interaction networks are structured across not only temporal, but also spatial or functional multiple layers. Species activity is likely to prove most advantageous given the strong emphasis to integrate multiple ecological processes, such as bellow-aboveground interactions, multitrophic interactions, or mutualistic-antagonistic networks (e.g., García-Callejas *et al*. 2018).

The most important bird and plant species in terms of species strength, number of links (i.e., degree), and multilayer centrality (i.e., versatility), were those with higher species activity. These birds (e.g., *Sylvia* spp., *E. rubecula*, and *T. merula*) are highly frugivorous during this period of the year and are among the most important avian seed dispersers in the Mediterranean basin (Herrera 2001). Therefore, their importance was not surprising as birds with a high degree of frugivory are fundamental to the structure of bird-fruit networks (Rumeu *et al*. 2017; Sebastián-González 2017). The birds with lower species activity and lower importance to the seed-dispersal network mainly include granivorous (e.g., *Chloris chloris*) and insectivorous species (e.g., *Cyanistes caeruleus*) that occasionally dispersed seeds from one or few plant species. As for plants, although the most important species in each year also tended to be those with higher species activity, one topologically important species in the seed-dispersal network was not temporally reliable: *Pistacia lentiscus* (Fig. 3). This species produces small and abundant lipid-rich fruits which are highly consumed by several bird species (Olesen *et al*. 2011a), but its fruits were only ripe during the study period in three of the years. Its absence in two networks was the result of a delay in the maturation of its fruits because it had unripe fruits during the sampling period. Nevertheless, when available, they were one of the most important fruit resources for birds. As some plants are known for highly variable fruit crops or fluctuations in the timing of fruit ripening (Herrera 1998), species activity may be a more accurate indicator of species importance for birds than for plants. Overall, these results indicate that bird species with high species activity tend to be more reliable seed dispersers. In turn, plants with high species activity provide reliable resources for several bird species, namely migrants that rely on fruits to increase their body fat reserves for the migratory flights (Bairlein 2002) Interestingly, there seems to be an independent relationship between species activity and plant and animal specialization d’ (Blüthgen *et al*. 2006). This is probably due to the generalist nature of most seed dispersers (Blüthgen *et al*. 2007), suggesting that the greater importance of temporally reliable species is not a reflection of a lower selectivity for resources.

Only six (50%) bird species and five (29%) plant species were present in all years; a similar relationship was observed in species turnover between years. These results contrast with previous findings of pollination studies interannual turnover which tends to be greater for animal than for plants (Petanidou *et al*. 2008; Dupont *et al*. 2009; Cirtwill *et al*. 2018). Part of this turnover might be related with the timing in fruit ripening of certain plant species (Herrera 1998), as already described here for *P. lentiscus*. The bird and plant species with higher species activity tend to dominate the network in terms of diversity and frequency of interactions. Consequently, the establishment of new interactions between temporally persisting species (i.e., rewiring) seems the main driver of interannual interaction turnover (e.g., Olesen *et al*. 2011b; CaraDonna *et al*. 2017). However, it is at this point difficult to distinguish true rewiring (i.e., new interactions taking place) from a normal undersampling of rare interactions in certain years (i.e., interactions that do occur but are not detected). Only nine links (12%) were observed on all years, indicating a low temporal link persistence. As observed in pollination studies (Chacoff *et al*. 2018), those few links were disproportionally frequent, suggesting that birds might predictably prefer to interact with the most temporally reliable plant species.

Our study revealed a relatively stable interannual network structure, with the noteworthy exception of 2014, when the network was not significantly nested. While the stability of seed-dispersal network structure across seasons has already been noted (Plein *et al*. 2013). Our study suggests that such stability can also be extended to interannual network structure, despite an appreciable species turnover, following the patterns observed in pollination networks (Alarcón *et al*. 2008; Petanidou *et al*. 2008; Dupont *et al*. 2009). However our study suggest that species roles tend to be temporally conserved in seed-dispersal networks, in contrast with pollination systems (Cirtwill *et al*. 2018). Our five-year study also shows that, as expected on any ecological process, not all years are exactly equal and that extrapolations based on temporally restricted sampling (such as nestedness in 2014) may lead to a biased characterization of network structure(Estes *et al*. 2018).

The identification of tight interaction modules within the relatively loose interaction networks has been one of the most insightful advances in community ecology of the last decade (e.g., Olesen *et al*. 2007; Schleuning *et al*. 2014) However, most ecological processes, including seed-dispersal, are continuous and not constrained by rigid temporal or spatial windows, likely affecting module detection. A multilayer modularly detection algorithm that is not constrained to seasonal or yearly data and where modules can span across multiple temporal or spatial layers, is likely to bring us much closer to the reality of natural communities (Mucha *et al*. 2010; Pilosof *et al*. 2017). This approach was used here for the first time to detect temporal seed-dispersal modules, which reveal to be very stable across years, with most species, especially birds, maintaining their module affiliation throughout the five consecutive years. This stability suggests that not only birds from different modules tend to have distinct fruit preferences, but these preferences tend to be temporally consistent and independent of fruit abundance. Indeed, some abundant plant species were rarely dispersed (e.g., *Arbutus unedo*), suggesting that birds likely select fruits based on other intrinsic traits such as their nutritional composition (Schaefer *et al*. 2003; Yang *et al*. 2013; Morán-López *et al*. 2018). Incorporating “historical interaction information” for module detection minimizes the influence of transient species roles and allows the detection of long-lasting modules which may be highly informative for conservation efforts (Blonder *et al*. 2012).

### Concluding remarks

Here, we implemented an innovative multilayer approach to understand the interannual dynamics of seed-dispersal networks and identified four temporally coherent interaction modules spanning across five consecutive years. The structure of the seed-dispersal network was relatively stable across years despite a significant turnover of species and interactions. Interestingly, the highly mobile birds, some of which migratory, presented a lower interannual turnover than their sessile mutualistic partners (i.e., fruiting plants). More importantly, our results revealed that species present across more years (defined here as having higher species activity) are also the most important on each year, both in terms of link richness and species strength, in a relationship independent from fruit availability and bird or plant specialization d’. Our results suggest that fruit-frugivore interactions are structured around a core of temporally reliable species, with which transient species tend to interact. By formally integrating species interacting on multiple spatial, temporal, or functional layers, multilayer networks are a most promising tool to approximate network analysis to the intrinsic complexity of natural communities.

## Acknowledgements

We thank to the Portuguese ringing authority (CEMPA/ICNF) for providing ringing permits and bird rings. This work was financed by FCT/MEC through national funds and co-funded by FEDER, within the PT2020 Partnership Agreement and COMPETE 2020 through grants UID/BIA/04004/2013, SFRH/BD/96292/2013 (J.M.C), SFRH/BD/77746/2011 (L.P.S.) and IF/00441/2013 (R.H.H.). R.H.H. was also supported by the Marie Curie action FP7-PEOPLE-2012-CIG-321794.

## References

Alarcón R., Waser N.M. & Ollerton J. (2008). Year-to-year variation in the topology of a plant-pollinator interaction network. Oikos.

Bairlein F. (2002). How to get fat: nutritional mechanisms of seasonal fat accumulation in migratory songbirds. Naturwissenschaften, 89, 1-10.

Barrat A., Barthelemy M., Pastor-Satorras R. & Vespignani A. (2004). The architecture of complex weighted networks. Proceedings of the National Academy of Sciences of the United States of America, 101, 3747-3752.

Bascompte J., Jordano P., Melián C.J. & Olesen J.M. (2003). The nested assembly of plant–animal mutualistic networks. PNAS, 100, 9383-9387.

Bates D., Maechler M., Bolker B. & Walker S. (2015). Fitting linear mixed-effects models using lme4. Journal of Statistical Software, 67, 1-48.

Blondel V.D., Jean-Loup G., Renaud L. & Etienne L. (2008). Fast unfolding of communities in large networks. Journal of Statistical Mechanics: Theory and Experiment, 2008, P10008.

Blonder B., Wey T.W., Dornhaus A., James R. & Sih A. (2012). Temporal dynamics and network analysis. Methods in Ecology and Evolution, 3, 958-972.

Blüthgen N., Menzel F. & Blüthgen N. (2006). Measuring specialization in species interaction networks. BCM Ecology, 6, 9.

Blüthgen N., Menzel F., Hovestadt T., Fiala B. & Blüthgen N. (2007). Specialization, constraints, and conflicting interests in mutualistic networks. Current Biology, 17, 341-346.

Brin S. & Page L. (2012). Reprint of: The anatomy of a large-scale hypertextual web search engine. Computer Networks, 56, 3825-3833.

Burkle L.A. & Alarcón R. (2011). The future of plant-pollinator diversity: understanding interaction networks across time, space, and global change. American Journal of Botany, 98, 528-538.

CaraDonna P.J., Petry W.K., Brennan R.M., Cunningham J.L., Bronstein J.L., Waser N.M. & Sanders N.J. (2017). Interaction rewiring and the rapid turnover of plant–pollinator networks. Ecology Letters, 20, 385-394.

Carnicer J., Jordano P. & Melián C.J. (2009). The temporal dynamics of resource use by frugivorous birds: a network approach. Ecology, 90, 1958-1970.

Chacoff N.P., Resasco J. & Vázquez D.P. (2018). Interaction frequency, network position, and the temporal persistence of interactions in a plant–pollinator network. Ecology, 99, 21-28.

Chao A. (1987). Estimating the population size for capture-recapture data with unequal catchability. Biometrics, 43, 783-791.

Cirtwill A.R., Roslin T., Rasmussen C., Olesen J.M. & Stouffer D.B. (2018). Between-year changes in community composition shape species’ roles in an Arctic plant–pollinator network. Oikos, 127, 1163-1176.

Colwell R.K. (2013). Estimates: Statistical estimation of species richness and shared species from samples. Version 9. User’s Guide and application published at: http://purl.oclc.org/estimates.

Correia M., Timόteo S., Rodríguez-Echeverría S., Mazars-Simon A. & Heleno R. (2016). Refaunation and the reinstatement of the seed-dispersal function in Gorongosa National Park. Conservation Biology, 31, 76-85.

Costa J.M., Ramos J.A., da Silva L.P., Timóteo S., Andrade P., Araújo P.M., Carneiro C., Correia E., Cortez P., Felgueiras M., Godinho C., Lopes R.J., Matos C., Norte A.C., Pereira P.F., Rosa A. & Heleno R.H. (2018). Rewiring of experimentally disturbed seed dispersal networks might lead to unexpected network configurations. Basic and Applied Ecology, 30, 11-22.

da Silva L.P., Ramos J.A., Coutinho A.P., Tenreiro P.Q. & Heleno R.H. (2016). Flower visitation by European birds offers the first evidence of interaction release in continents. Journal of Biogeography, 44, 687-695.

De Domenico M., Porter M.A. & Arenas A. (2015a). MuxViz: a tool for multilayer analysis and visualization of networks. Journal of Complex Networks, 3, 159-176.

De Domenico M., Solé-Ribalta A., Omodei E., Gómez S. & Arenas A. (2015b). Ranking in interconnected multilayer networks reveals versatile nodes. Nature Communications, 6, 6868.

Donatti C.I., Guimarães P.R., Galetti M., Pizo M.A., Marquitti F.M.D. & Dirzo R. (2011). Analysis of a hyper-diverse seed dispersal network: modularity and underlying mechanisms. Ecology Letters, 14, 773-781.

Dormann C.F., Gruber B. & Fründ J. (2008). Introducing the bipartite Package: Analysing Ecological Networks. R News, 8, 8-11.

Dupont Y.L., Padrón B., Olesen J.M. & Petanidou T. (2009). Spatio-temporal variation in the structure of pollination networks. Oikos, 118, 1261-1269.

Estes L., Elsen P.R., Treuer T., Ahmed L., Caylor K., Chang J., Choi J.J. & Ellis E.C. (2018). The spatial and temporal domains of modern ecology. Nature Ecology & Evolution, 2, 819-816.

Galeano J., Pastor J.M. & Iriondo J.M. (2009). Weighted-Interaction Nestedness Estimator (WINE): A new estimator to calculate over frequency matrices. Environmental Modelling & Software, 24, 1342-1346.

García-Callejas D., Molowny-Horas R. & Araújo M.B. (2018). Multiple interactions networks: towards more realistic descriptions of the web of life. Oikos, 127, 5-22.

González-Castro A., Yang S., Nogales M. & Carlo T.A. (2012). What determines the temporal changes of species degree and strength in an oceanic island plant-disperser network? PLoS ONE, 7, e41385.

Heleno R.H., Olesen J.M., Nogales M., Vargas P. & Traveset A. (2013). Seed dispersal networks in the Galápagos and the consequences of alien plant invasions. Proceedings of the Royal Society B: Biological Sciences, 280.

Herrera C.M. (1984). A study of avian frugivores, bird-dispersed plants, and their interaction in Mediterranean scrublands. Ecological Monographs, 54, 1-23.

Herrera C.M. (1998). Long-term dynamics of Mediterranean frugivorous birds and fleshy fruits: a 12-years study. Ecological Monographs, 68, 511-538.

Herrera C.M. (2001). Dispersión de semillas por animales en el Mediterráneo: ecología y evolución. In: Ecosistemas Mediterráneos (eds. Rodriguez RZ & Iraola FIP). Servicio Publicaciones CSIC Madrid, pp. 125-152.

Jordano P. (1987). Patterns of mutualistic interactions in pollination and seed dispersal: connectance, dependence asymmetries, and coevolution. The American Naturalist, 129, 657-677.

Jordano P. (2014). Fruits and frugivory. In: Seeds: the ecology of regeneration in plant communities (ed. Gallagher RS). CABI Wallingford, U.K., pp. 18-61.

Jordano P., García C., Godoy J.A. & García-Castaño J.L. (2007). Differential contribution of frugivores to complex seed dispersal patterns. Proceedings of the National Academy of Sciences, 104, 3278-3282.

Jutla I.S., Jeub L.G.S. & Mucha P.J. (2014). A generalized Louvain method for community detection implemented in MATLAB, http://netwiki.amath.unc.edu/GenLouvain (20112014). In.

Koleff P., Gaston K.J. & Lennon J.J. (2003). Measuring beta diversity for presence–absence data. Journal of Animal Ecology, 72, 367-382.

McLeod A.I. (2011). Kendall: Kendall rank correlation and Mann-Kendall trend test. R package version 2.2. https://CRAN.R-project.org/package=Kendall.

Morán-López T., Carlo T.A., Amico G. & Morales J.M. (2018). Diet complementation as a frequency-dependent mechanism conferring advantages to rare plants via dispersal. Functional Ecology, 0.

Mucha P.J., Richardson T., Macon K., Porter M.A. & Onnela J.-P. (2010). Community structure in time-dependent, multiscale, and multiplex networks. Science, 328, 876-878.

Newman M.E.J. (2001). Scientific collaboration networks. II. Shortest paths, weighted networks, and centrality. Physical Review E, 64, 016132.

Nicosia V. & Latora V. (2015). Measuring and modeling correlations in multiplex networks. Physical Review E, 92, 032805.

Oksanen J., Blanchet F.G., Kindt R., Legendre P., Minchin P.R., O’Hara R.B., Simpson G.L., Solymos P., Stevens M.H.H. & Wagner H. (2015). vegan: Community Ecology Package. R package version 2.2-1. In.

Olesen J.M., Bascompte J., Dupont Y.L., Elberling H., Rasmussen C. & Jordano P. (2011a). Missing and forbidden links in mutualistic networks. Proc. R. Soc. B, 278, 725-732.

Olesen J.M., Bascompte J., Dupont Y.L. & Jordano P. (2007). The modularity of pollination networks. PNAS, 104, 19891-19896.

Olesen J.M., Dupont Y.L., O’Gorman E., Ings T.C., Layer K., Melián C.J., Trøjelsgaard K., Pichler D.E., Rasmussen C. & Woodward G. (2010). From Broadstone to Zackenberg: space, time and hierarchies in ecological networks. Advances in Ecological Research, 42, 1-69.

Olesen J.M., Stefanescu C. & Traveset A. (2011b). Strong, long-term temporal dynamics of an ecological network. PLoS ONE, 6, e26455.

Opsahl T. (2009). Structure and Evolution of Weighted Networks. University of London (Queen Mary College), London, UK, pp. 104-122.

Opsahl T. (2013). Triadic closure in two-mode networks: Redefining the global and local clustering coefficients. Social Networks, 35, 159-167.

Patefield W.M. (1981). Algorithm AS 159: An efficient method of generating random R × C tables with given row and column totals. Journal of the Royal Statistical Society. Series C (Applied Statistics), 30, 91-97.

Petanidou T., Kallimanis A.S., Tzanopoulos J., Sgardelis S.P. & Pantis J.D. (2008). Long-term observation of a pollination network: fluctuation in species and interactions, relative invariance of network structure and implications for estimates of specialization. Ecology Letters, 11, 564-575.

Pilosof S., Porter M.A., Pascual M. & Kéfi S. (2017). The multilayer nature of ecological networks. Nature Ecology & Evolution, 1, 0101.

Plein M., Längsfeld L., Neuschulz E.L., Schultheiß C., Ingmann L., Töpfer T., Böhning-Gaese K. & Schleuning M. (2013). Constant properties of plant–frugivore networks despite fluctuations in fruit and bird communities in space and time. Ecology, 94, 1296-1306.

Poisot T. (2016). betalink: Beta-Diversity of Species Interactions. R package version 2.2.1. https://CRAN.R-project.org/package=betalink.

Poisot T., Canard E., Mouillot D., Mouquet N. & Gravel D. (2012). The dissimilarity of species interaction networks. Ecology Letters, 15, 1353-1361.

R Core Team (2017). R: A language and environment for statistical computing. R Foundation for Statistical Computing, Vienna, Austria. URL http://www.R-proiect.org/.

Ramos-Robles M., Andresen E. & Díaz-Castelazo C. (2016). Temporal changes in the structure of a plant-frugivore network are influenced by bird migration and fruit availability. PeerJ, 4, e2048.

Rumeu B., Devoto M., Traveset A., Olesen J.M., Vargas P., Nogales M. & Heleno R. (2017). Predicting the consequences of disperser extinction: richness matters the most when abundance is low. Functional Ecology, 31, 1910-1920.

Schaefer H.M., Schmidt V. & Bairlein F. (2003). Discrimination abilities for nutrients: which difference matters for choosy birds and why? Animal Behaviour, 65, 531-541.

Schleuning M., Ingmann L., Strauß R., Fritz S.A., Dalsgaard B., Matthias Dehling D., Plein M., Saavedra F., Sandel B., Svenning J.-C., Böhning-Gaese K. & Dormann C.F. (2014). Ecological, historical and evolutionary determinants of modularity in weighted seed-dispersal networks. Ecology Letters, 17, 454-463.

Sebastián-González E. (2017). Drivers of species’ role in avian seed-dispersal mutualistic networks. Journal of Animal Ecology, 86, 878-887.

Tilman D., Isbell F. & Cowles J.M. (2014). Biodiversity and ecosystem functioning. Annual Review of Ecology, Evolution, and Systematics, 45, 471-493.

Timόteo S., Correia M., Rodríguez-Echeverría S., Freitas H. & Heleno R. (2018). Multilayer networks reveal the spatial structure of seed-dispersal interactions across the Great Rift landscapes. Nature Communications, 9, 140.

Traveset A., Heleno R.H. & Nogales M. (2014). The ecology of seed dispersal. In: Seeds: the ecology of regeneration in plant communities (ed. Gallagher RS). CABI Wallingford, pp. 62-93.

Trøjelsgaard K., Jordano P., Carstensen D.W. & Olesen J.M. (2015). Geographical variation in mutualistic networks: similarity, turnover and partner fidelity. 282.

Trøjelsgaard K. & Olesen J.M. (2016). Ecological networks in motion: micro- and macroscopic variability across scales. Functional Ecology, 30, 1926-1935.

Vázquez D.P., Morris W.F. & Jordano P. (2005). Interaction frequency as a surrogate for the total effect of animal mutualists on plants. Ecology Letters, 8, 1088-1094.

Wenny D.G., Şekercioğlu Ç., Cordeiro N.J., Rogers H.S. & Kelly D. (2016). Seed dispersal by fruit-eating birds. In: Why birds matter: avian ecological function and ecosystem services (eds. Şekercioğlu Ç, Wenny DG & Whelan CJ). The University of Chicago Press, pp. 107-145.

Whittaker R.H. (1960). Vegetation of the Siskiyou mountains, Oregon and California. Ecological Monographs, 30, 279-338.

Yang S., Albert R. & Carlo T.A. (2013). Transience and constancy of interactions in a plant-frugivore network. Ecosphere, 4, 147.

